# Goslin - A Grammar of Succinct Lipid Nomenclature

**DOI:** 10.1101/2020.04.17.046656

**Authors:** Dominik Kopczynski, Nils Hoffmann, Bing Peng, Robert Ahrends

## Abstract

We introduce Goslin, a polyglot grammar for common lipid shorthand nomenclatures based on the LipidMaps nomenclature and the shorthand nomenclature established by Liebisch *et al*. and used by LipidHome and SwissLipids. Goslin was designed to address the following pressing issues in the lipidomics field: 1) to simplify the implementation of lipid name handling for developers of mass spectrometry-based lipidomics tools; 2) to offer a tool that unifies and normalizes the main existing lipid name dialects enabling a lipidomics analysis in a high-throughput fashion. We provide implementations of Goslin in four major programming languages, namely C++, Java, Python 3, and R to kick-start adoption and integration. Further, we set up a web service for users to work with Goslin directly. All implementations are available free of charge under a permissive open source license.

## Introduction

Lipids are beside DNA/RNA, proteins, and metabolites the most frequently occurring biomolecules in organic cells. Several lipid classes have a similar molecular structure. Therefore, lipid names were designed to represent their systematic structure in a clear and concise way to help classify and therefor distinguish lipids by name.(1, 2) Within the past two decades, a systematic shorthand notation for lipids has evolved, spurred by initial standardization approaches within the LipidMaps consortium.(3, 4) With the advent of high-resolution mass spectrometry (MS), as the key tool to investigate lipids, the hierarchical representation of lipids was extended to be able to report multiple levels of structural knowledge.(5, 6) Since mass spectrometry is currently the dominant technology for the identification and quantification of lipids,(7, 8) these notations are primarily designed to satisfy the requirements in that application area. Structural knowledge about lipids identified with MS can be represented as a hierarchical tree or table (see Table 1), as introduced by LipidHome(9) and later refined by SwissLipids(10). This tree is rooted at category level (e.g., glycerophospholipids), and descends from class (e.g., glycerophosphoethanolamine) to species (e.g., phosphatidylethanolamine(32:1), PE(32:1)), to molecular subspecies (e.g. PE(16:0_16:1) with determined fatty acyl (FA) lengths and saturation), then to structural subspecies (e.g. PE(16:1/16:0) with defined stereospecific numbering (SN) positions for the FAs), and ultimately to isomeric level (e.g. PE(16:1(6Z)/16:0) or 1-(6Z-hexadecenoyl)-2-hexadecanoyl-sn-glycero-3-phosphoethanolamine). Using the shorthand abbreviations for the lipid classes (e.g., *PE* for phosphatidylethanolamine or *SM* for sphingomyelin together with the shorthand notation for various variable building blocks, such as length of fatty acyl chains and long chain bases, as well as the number of double bonds and hydroxylations, lipid names become both manageable and descriptive. With the current developments in lipidomics and the upcoming high-throughput analyses, an increasing demand for automated computational data handling is on the horizon. Hence, it is absolutely essential that identified lipids are correctly and unambiguously named and stored for follow-up analyses during a computational lipid analysis workflow that may consist of several consecutive tools. Lipid naming has however evolved into several dialects which complicates the unified computational treatment and parsing of lipid names. For instance, in literature one may find for essentially the same lipid species any of the following names:

Cer(d18:1/16:0),
Cer(d18:1/16:0),
d18:1/16:0 Cer,
Cer 18:1;2/16:0, or
CER[N(16)S(18)].

**Table 1.** Hierarchical classification of PE(16:1(6Z)/16:0). Higher levels represent the lipid with less detail and more summarized information, e.g. the top most category level reports that PE(32:1) as a species is a Glycerophospholipid. The species level expresses only the head group (HG), and the sum of # carbon atoms and # double bonds in both fatty acyls (FAs). Intermediate levels report additional details about FA composition (molecular subspecies) and SN position with respect to the head group (structural subspecies). At the lowest level, the FA composition (# carbon atoms, # double bonds), their SN position (first is SN1, second is SN2), and the position of the double bond are expressed (6th carbon in SN1 FA, in Z configuration). The ’Level’ column details the mapping of Goslin’s levels to the respective levels in the LipidMaps (LM) and SwissLipids hierarchies (SL). Hydroxyl groups have been omitted from this example for brevity, but they are also fully supported by Goslin.

| Level | Name | Description |
| --- | --- | --- |
| Category (LM) | Glycerophospholipids | Lipid category |
| Class (LM) | Glycerophosphoethanolamine | Lipid class |
| Species (SL, LM Subclass) | Phosphatidylethanolamine (32:1), PE(32:1) | HG, FA summary |
| Molecular Subspecies (SL) | PE(16:0_16:1) | HG, two FAs |
| Structural Subspecies (SL) | PE(16:1/16:0) | HG, SN pos. for FA1 and FA2 |
| Isomeric Subspecies (SL, LM) | PE(16:1(6Z)/16:0) | HG, SN pos., Stereo |

Although some of these lipid names may look very similar or differ perhaps only in one character, a simple string matching fails when these names are not exactly equal. As a consequence, long and error-prone manual curation is necessary in order to streamline lists of lipid names for their processing in follow-up analysis scripts, workflows, or tools, or for their submission to research data repositories.

## Results

To overcome these challenges, we developed the ’Grammar of Succinct Lipid Nomenclature’ (*Goslin*). It is designed to act as a library for the development of lipidomics tools providing a standardized data structure for storing structural lipid information. The parsing of lipid names as well as the lipid name generation are the main functions of Goslin. We therefor defined a context free grammar(11) that defines rules and productions for all structural properties of the lipid nomenclature, including mass spectrometry specific information about unlabeled and heavy isotope labeled species, as well as fragments. A short extract of that grammar is illustrated in Figure 1. Currently, the grammar covers 289 lipid classes within the seven most occurring lipid categories in eukaryotic organisms, namely fatty acyls, glycerolipids, glycerophospholipids, saccharolipids, sphingolipids, sterol lipids, and polyketides (for a detailed lipid list, see Supplementary Table 1). To keep all implementations up to date, one authoritative lipid list containing the current lipid name abbreviation including all its synonyms is used by the different implementations. The major advantages of using a grammar rather than a manually coded parser are its flexibility and extensibility. Regular expressions are also not suitable for parsing lipid names, since they are incapable of recognizing nested patterns and can only recognize words from regular languages.(11, 12) Obviously, lipid names do not exhibit such a regular structure. Changes or updates to the lipid nomenclature can easily be applied to the grammar. Parsers based on these grammars can be generated automatically(13) to check whether a string is a valid word within a language defined by the grammar. Goslin is able to map lipid names to either species level (which includes category and class), subspecies level, to molecular and structural subspecies level, as well as to isomeric subspecies level. Another advantage is its compatibility with lipid nomenclatures from existing publications. For instance, we defined additional grammars capable of parsing the structural lipid names from LipidMaps(3, 4), SwissLipids(10), and HMDB.(14) Variations (e.g., a blank after the head group abbreviation or none) are already defined within the grammar and handled automatically without the need for manually coded handling or additional validation. We also consider and handle different common abbreviations of the head groups (e.g., SPH vs. Sph or DG vs. DAG) and support different strings as separator symbols between multiple fatty acyl chains (either ’-’, ’_’, or ’/’ from literature). Changes of these symbols can easily be updated in the grammar. Given a lipid name on a certain hierarchy level, the Goslin parser implementation is also able to report the lipid names for all of its parent levels. This feature simplifies several use-cases, such as the computation of the distribution of lipids among the lipid classes or lipid categories. The polyglot approach already covers several existing dialects of the lipid nomenclature and can be extended arbitrarily to cover new lipid categories and classes, head groups and FA modifications. Tailored event handlers (one for each grammar) transfer all recognized elements of the structural lipid name into a common data structure that reflects the hierarchy similarly to that used in LipidHome(9) and SwissLipids.(10) Our implementations are written in C++ (version 11 and higher), Java (version 11 and higher), Python (version 3), and R (version 3.6 and higher) and can easily be included into existing lipidomics tools. Detailed descriptions on the integration and usage of the programming libraries are provided in the Supplementary Information. Additionally, we provide several instructive code snippets on the usage of Goslin for input and output of lipid names. We benchmarked our implementations with respect to their execution time for parsing. As a benchmark device, we used an Intel(R) Xeon(R) 2.80 GHz quad core desktop computer with 16GB RAM. The results are listed in Table 2. All implementations are fast enough to parse lipid name lists of regular experiment sizes (about 1000 lipid names) within a second. To provide a user-friendly alternative to the libraries and command line interface (CLI), we developed a Goslin web application (see https://apps.lifs.isas.de/goslin) that offers form-based lipid name translation, as well as a REST API for language agnostic, programmatic translation.

**Figure 1.**
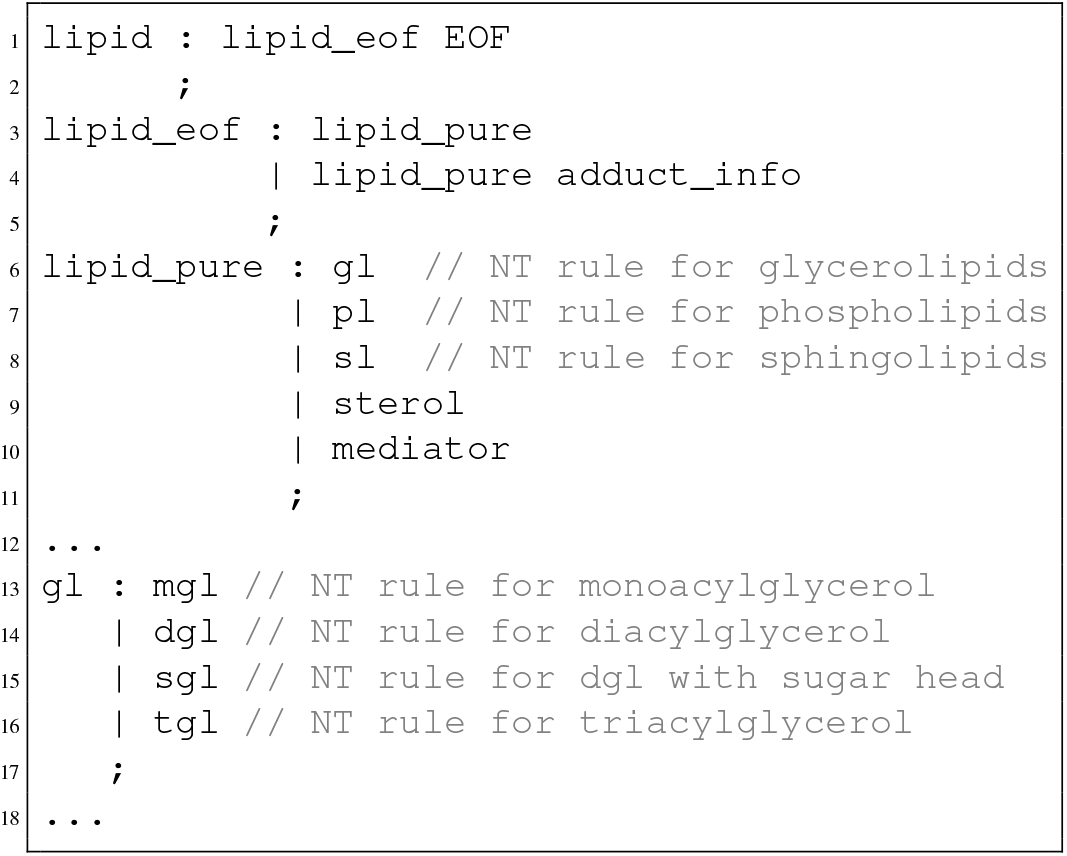
Snippet of the Goslin grammar: on the left hand side of each colon, one non-terminal (*NT*) rule defines a set of production rules which are either terminal or non-terminal rules. For instance, the production rule `lipid_eof` defines a lipid name by identifying either a pure lipid name or a pure lipid name followed by an identified scheme of an adduct. The complete grammars are provided in our repositories (see https://github.com/lifs-tools/goslin).

**Table 2.** Time performance benchmark on the different grammars with different lipid name test sets. All values for the programming languages C++, Java, and Python 3 have the unit *parsed lipids per second*. All testsets contain 10000 lipid names parsable by their grammar. Here, the testset Goslin (S) contains rather short lipid names on average and Goslin (L) rather long lipid names. The parsing time is dependent on the length of the lipid string. Therefore, the average length (*avg len*) over all lipids in each data set is reported. Note, that the R implemenation utilizes the C++ library internally. Therefore, it is not listed here explicitly.

| Test set | avg len | C++ | Java | Python |
| --- | --- | --- | --- | --- |
| Goslin (S) | 15.59 | 8237.23 | 7047.57 | 2957.81 |
| Goslin (L) | 37.76 | 2880.18 | 7771.71 | 807.31 |
| LipidMaps | 38.57 | 3727.17 | 5498.62 | 996.36 |
| SwissLipids | 36.84 | 3691.4 | 5810.29 | 973.80 |
| Hmdb | 32.41 | 4266.21 | 9100.97 | 1280.73 |

Lipid names can be directly copied and pasted into the submission form of the web application or can be uploaded as a file, with one lipid name per line. Upon submission, the application parses all lipid names and provides a result list with all identified lipids and additional structural information, as well as cross-links to LipidMaps and SwissLipids, where applicable. For lipid names that can not be parsed, it reports specific errors within the submission form to help the user fix the name. The result list can also be downloaded in a spreadsheet friendly tab-separated value format. For programmatic access, the user can send a web request (HTTPS) to translate her lipid list using the provided REST application programming interface (API) by sending a POST request with a JavaScript Object Notation (JSON) body (JSON list of lipid name strings). The response is a JSON list with embedded objects (for usage and tutorials, see Supplementary Information). The REST API is meant to be used by automated workflows, when neither of our offline implementations is suitable or applicable. We also offer instructions to build a Docker image of the web-application for local and customized deployments, as well as a Bioconda recipe for the jgoslin CLI(15) for inclusion in automated bioinformatics workflows.

## Conclusions

With Goslin, we provide another building block towards a more harmonized and inter-operable bioinformatics tool landscape for lipidomics. Equipped with multiple grammars, Goslin is able to interpret several lipid name dialects, such as the LipidMaps nomenclature, the shorthand nomenclature by Liebisch *et al*., the SwissLipids nomenclature, and the HMDB nomenclature. Goslin translates them into a normalized, up-to-date nomenclature and structured representation. Currently, we are working on several improvements of the library, 1) to offer an implementation of Goslin in additional programming languages (e.g., C#); 2) to improve the performance of the existing implementations; and 3) to add more structural lipid classes and categories into the grammar for a higher coverage of lipids. We already applied Goslin in other recent projects(16, 17) and achieved a high performance boost especially when exchanging our lipid lists with our collaboration partners. The Goslin implementations are freely available under https://github.com/lifs-tools/goslin under the terms of liberal open source licenses.

## Author contributions

D.K., N.H. designed the grammars, N.H. implemented the Java and R libraries and the Goslin web-application. D.K. implemented the C++ and Python versions. B.P. and R.A. contributed their structural and chemical knowledge of lipids. R.A. supervised the project. All authors wrote and approved of the manuscript.

## ACKNOWLEDGEMENTS

The supports by the Ministerium für Kultur und Wissenschaft des Landes Nordrhein-Westfalen and the Regierende Bürgermeister von Berlin, Senatskanzlei Wissenschaft und Forschung - inkl. Wissenschaft und Forschung, and the Bundesministerium für Bildung und Forschung (BMBF) to DK (Code 031A534B), NH (Code 031L0108A) and RA are gratefully acknowledged. We thank Harald Köfeler and Gerhard Liebisch for the close collaboration within the International Lipidomics Society on lipidomics standardization.

