## Supplementary Information for "Goslin - A Grammar of Succinct Lipid Nomenclature"

#### Contents

**1 Web Application and REST API** **2**

**2 C++ Implementation** **9**

**3 Python Implementation** **12**

**4 R Implementation** **16**

**5 Java Implementation** **19**

**6 Goslin Object Model** **23**

**7 List of Supported Lipids** **24**

### 1 Web Application and REST API

#### Interactive Usage

The interactive grammar of succinct lipid nomenclatures (Goslin) web application is available at `https://apps.lifs.isas.de/goslin`. It provides two forms to i) upload a file containing one lipid name per line (see Supplementary Figure 1), or ii) upload a list of lipid names, defined by the user in an interactive form (see Supplementary Figure 2). The latter form also allows pasting lists of lipid names directly from the clipboard with `CTRL+V`. Both forms provide feedback for issues concerning every processed lipid, such as invalid names or typos (see Supplementary Figure 3), to allow the user to cross-check their data before proceeding.

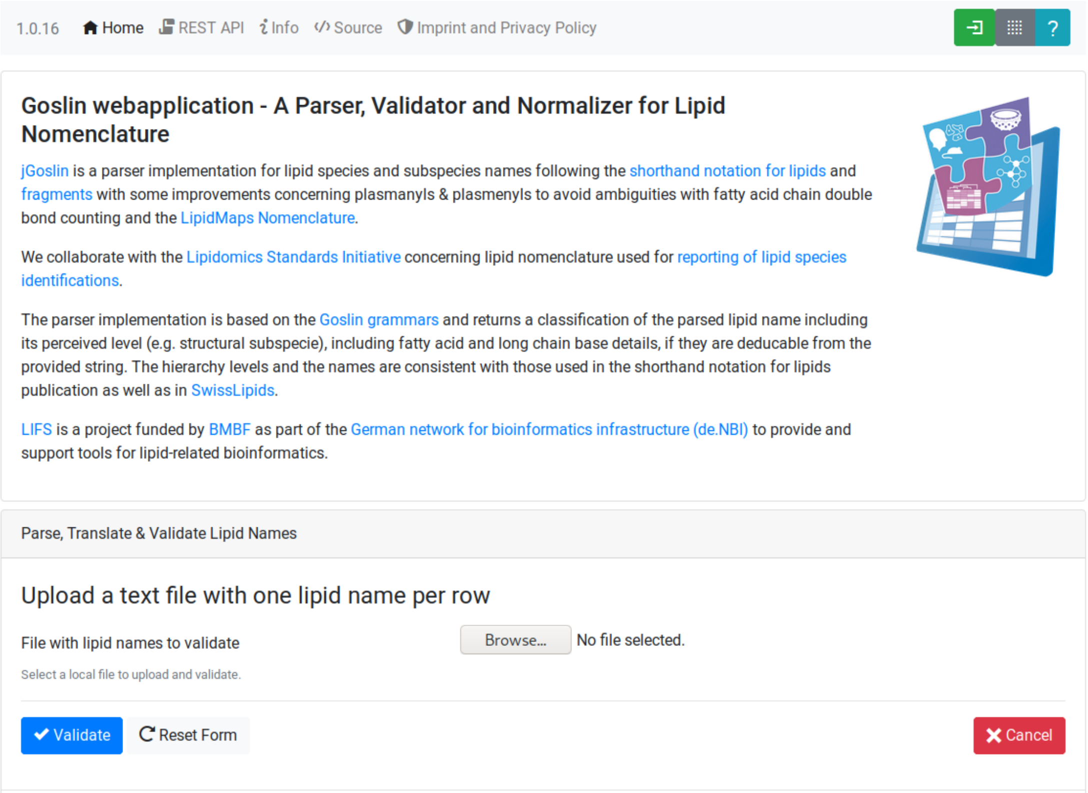


Supplementary Figure 1: Goslin web application submission form for text files with one lipid name per row.

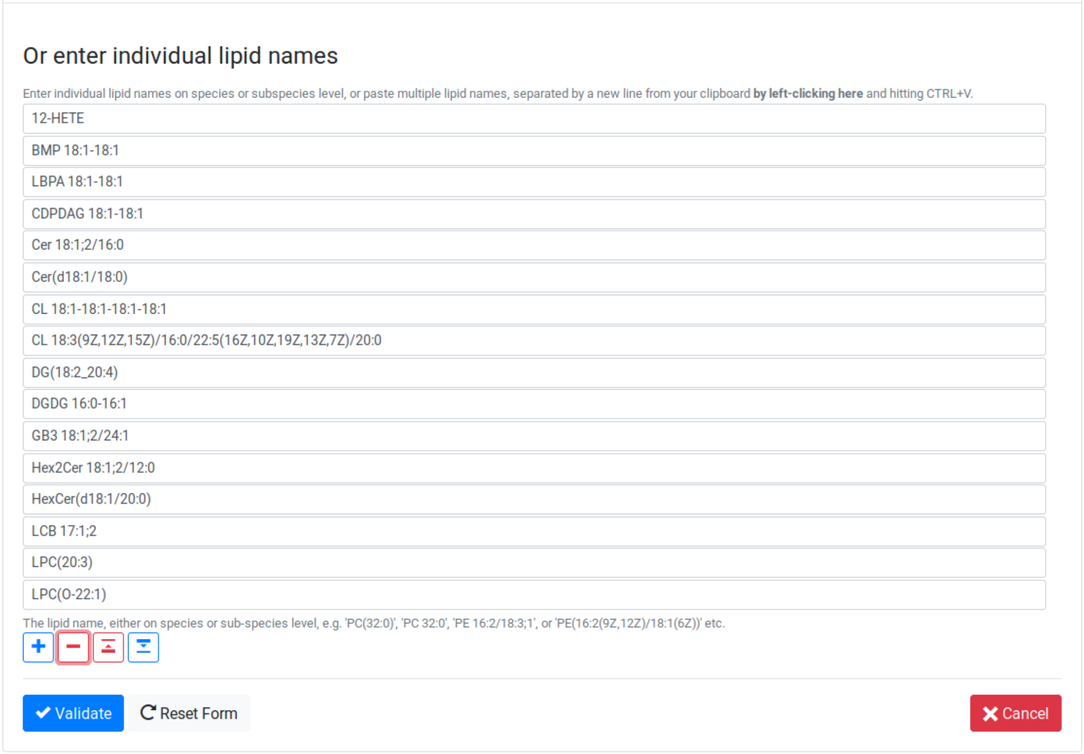


Supplementary Figure 2: Goslin web application submission form for user-defined lipid names.

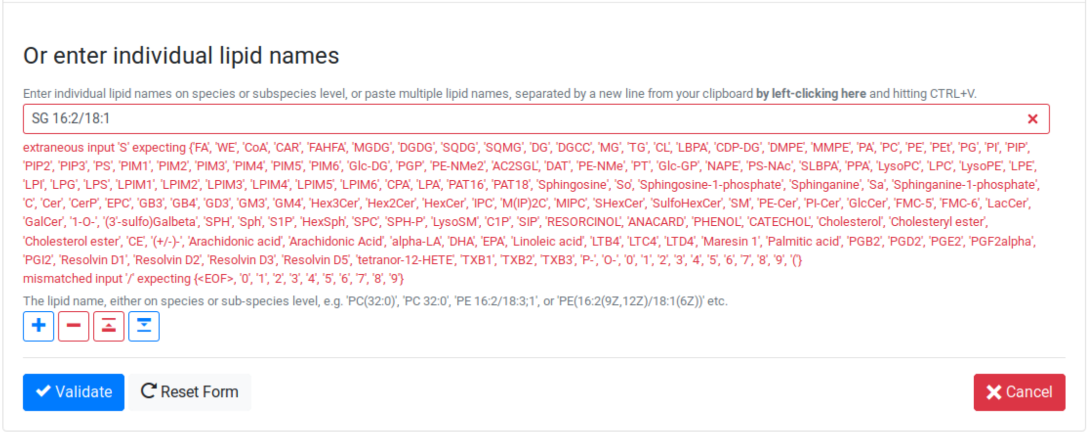


Supplementary Figure 3: Goslin web application submission form for user-defined lipid names provides feedback for unknown or unsupported names and parts thereof.

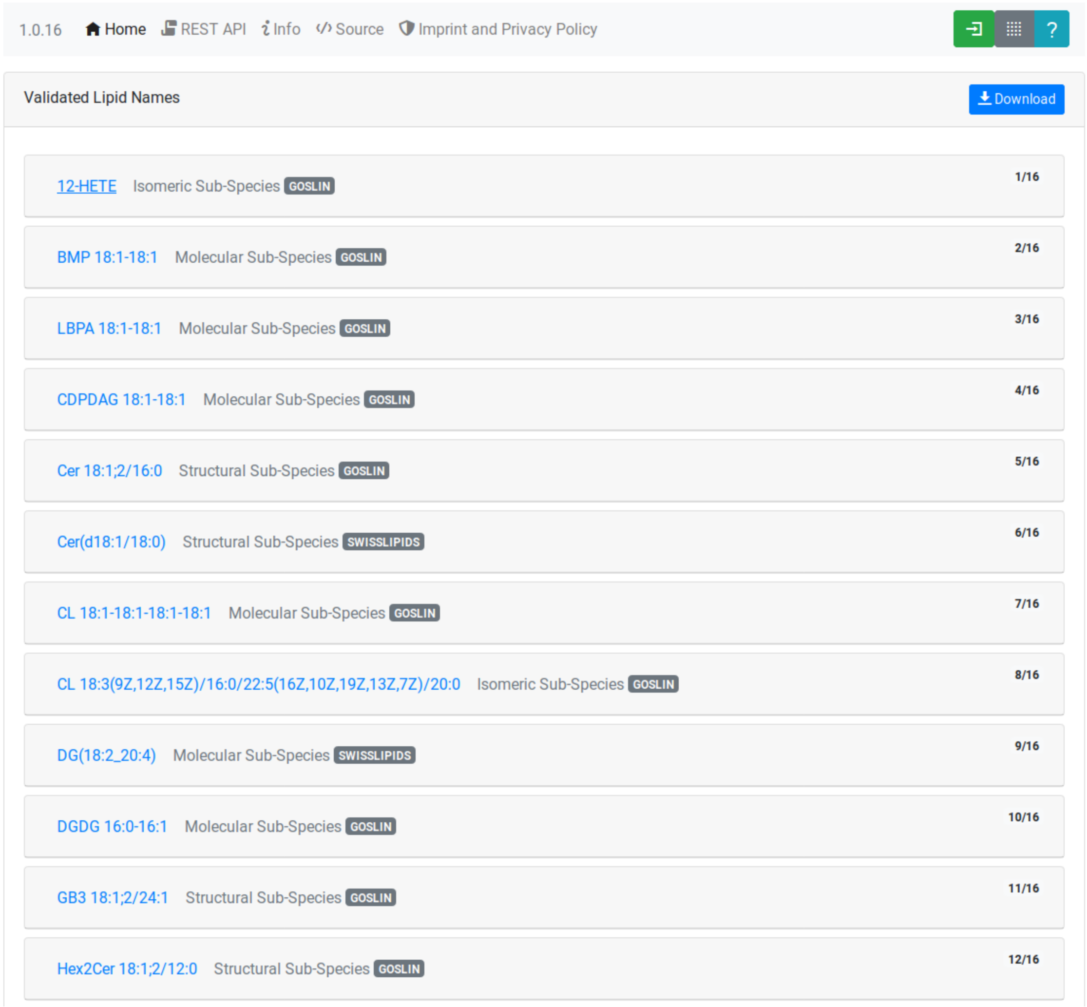


Supplementary Figure 4: Parsing results are displayed as 'cards' for every lipid name. Clicking on a card opens it and shows details of the according lipid.

After successful validation, the validated lipids are returned in overview cards (see Supplementary Figure 4), detailing their LipidMAPS classification[1], cross-links to SwissLipids[2] and/or LipidMAPS or HMDB[3]. Additionally, the cards show summary information about the number of carbon atoms, double bonds, hydroxylations and detailed information, such as double bond position, long-chain-base status, and the bond type of the fatty acyl to the head group for each fatty acyl, if available (see Supplementary Figure 5) .

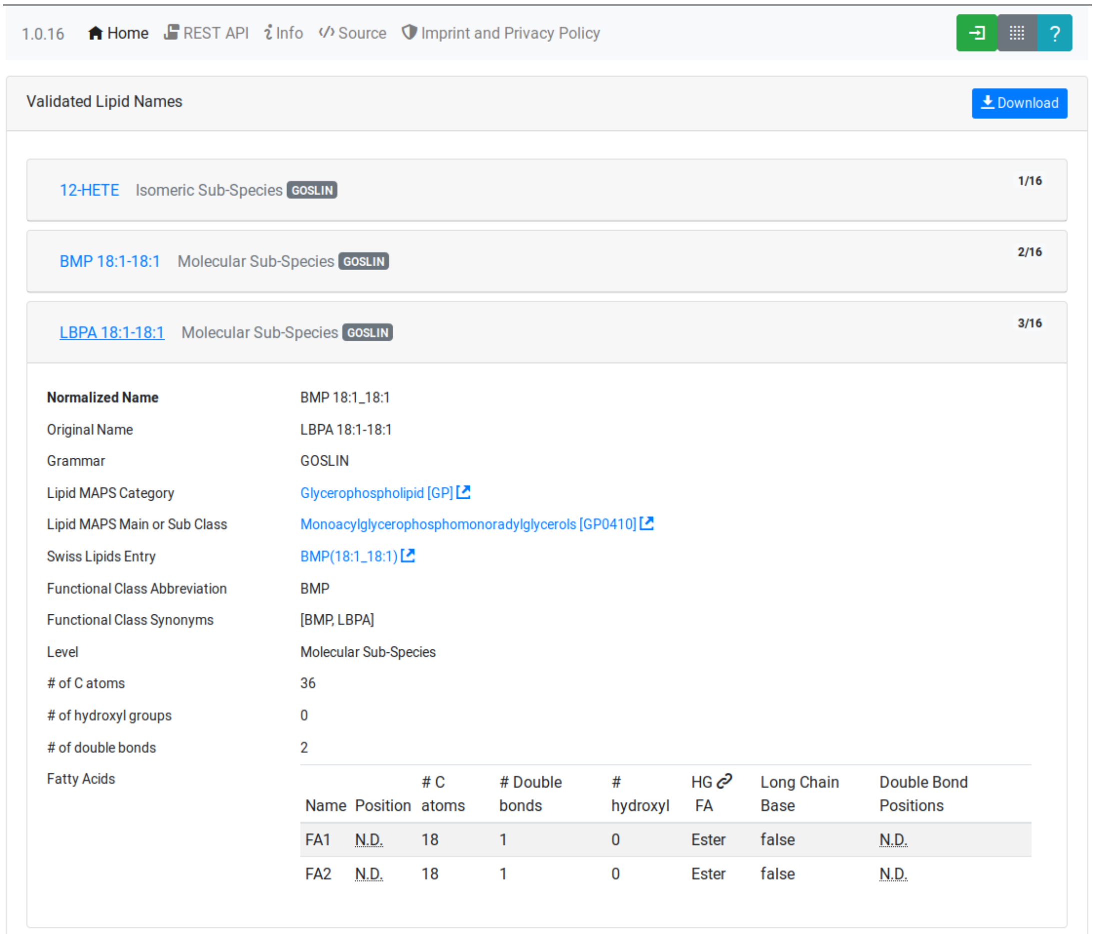


Supplementary Figure 5: Each result card displays summary and detail information about a lipid. Depending on the lipid level, this can include information about each individual fatty acyl. Cross-links to SwissLipids and LipidMAPS are shown where a normalized lipid name could be matched unambiguously against the normalized names of SwissLipids and / or LipidMAPS lipids.

The source code for the web application and instructions to build it as a Docker container are available at `https://github.com/lifs-tools/goslin-webapp` under the terms of the open source Apache license version 2.

#### Programmatic access via the REST API

An interactive documentation for the representational state transfer (REST) application programming interface (API) of the Goslin web application is available at `https://apps.lifs.isas.de/goslin/swagger-ui.html` (see Supplementary Figure 6). To illustrate its usage, we will briefly show a small example how a user can access the REST API with a standard hypertext transfer protocol (HTTP) client.

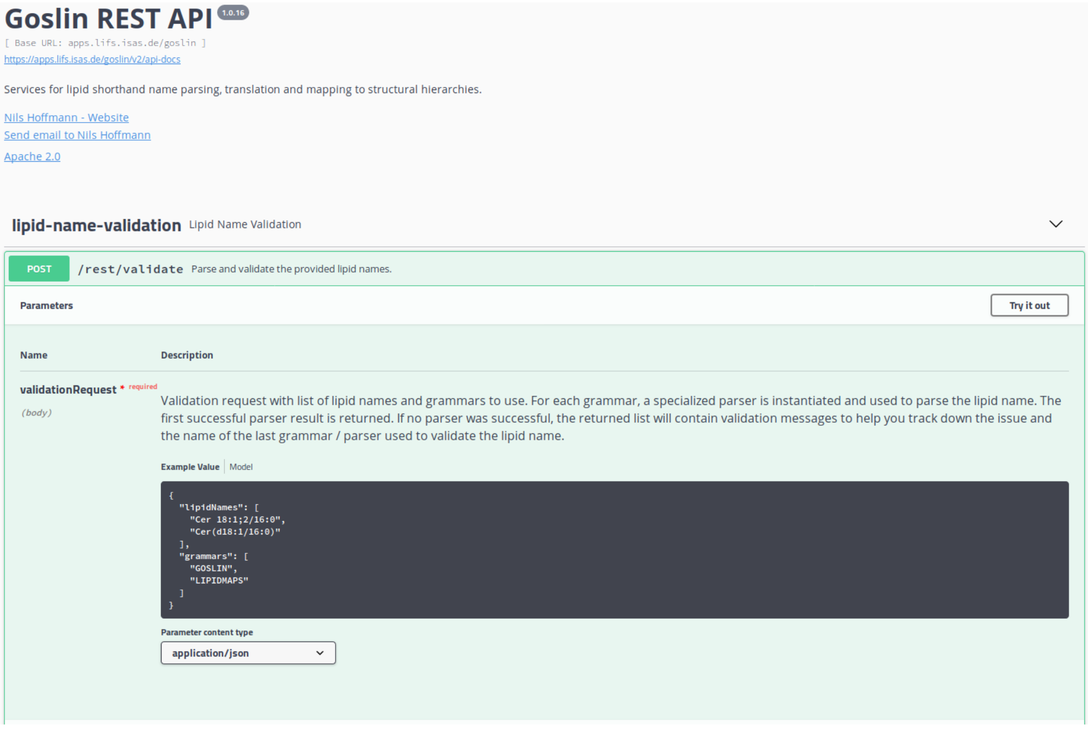


Supplementary Figure 6: The Goslin web application provides an interactive documentation for its REST API to simplify programmatic access.

The Structure for the request consists of a JavaScript object notation (JSON) object {} enclosing two lists, with the names `lipidNames` and `grammars`. Acceptable values for `grammars` are: `LIPIDMAPS`, `GOSLIN`, `GOSLIN_FRAGMENTS`, `SWISSLIPIDS`, and `HMDB`. A complete list is available from the interactive REST API documentation's `Models` section under `ValidationRequest`. Both fields in the `ValidationRequest` accept comma-separated entries, enclosed in double quotes:

```
{
    "lipidNames": [
      "Cer(d18:1/16:1(6Z))"
    ],
    "grammars": [
      "LIPIDMAPS"
    ]
}
```

Sending the HTTP POST request with `curl` as an HTTP client looks as follows:

```
curl -X POST "https://apps.lifs.isas.de/goslin/rest/validate" -H "accept: */*" -H
```

```
"Content-Type: application/json" -d "{ \"lipidNames\": [
\"Cer(d18:1/16:1(6Z))\" ], \"grammars\": [ \"LIPIDMAPS\" ]}"
```

The REST API will return the following result for the request, with a HTTP response code of 200 (OK). This result returns a map of properties for each lipid name that was parsed. If at least one name is not parseable, the REST API will return a response code of 400 (Client error), together with the same results reponse object. In that case, the `failedToParse` field in the response will contain the number of lipid names that could not be parsed. For those results where no grammar was applicable, the `grammar` field will contain the string `NOT_PARSEABLE`.¸In other cases, that field will contain the last grammar used to parse the lipid name and the `messages` field will contain a list of validation messages that help to narrow down the offending bits in the lipid name.

```
{
  "results": [
    {
      "lipidName": "Cer(d18:1/16:1(6Z))",
      "grammar": "LIPIDMAPS",
      "messages": [],
      "lipidAdduct": {
        "lipid": {
          "lipidCategory": "SP",
          "lipidClass": "CER",
          "headGroup": "Cer",
          "info": {
            "type": "STRUCTURAL",
            "name": "Cer",
            "position": -1,
            "lipidFaBondType": "ESTER",
            "lcb": false,
            "modifications": [],
            "doubleBondPositions": {},
            "level": "STRUCTURAL_SUBSPECIES",
            "ncarbon": 34,
            "nhydroxy": 2,
            "ndoubleBonds": 2
          },
```

The response part also reports the normalized name (`goslinName`), as well as classification information using the LipidMAPS category and class associated to the parsed lipid.

```
      },
      "goslinName": "Cer 18:1;2/16:1(6Z)",
      "lipidMapsCategory": "SP",
      "lipidMapsClass": "SP0203",
```

The response also reports information on the fatty acyls detected in the lipid name. In this case, a long chain base (LCB) (in the ceramide) has been detected. The name given here as an example was classified on structural subspecies level, since the LCB contains one double bond, but without positional E/Z information. The fatty acyl FA1 at the sn2 position does report E/Z information for its double bond, thus FA1 is an isomeric fatty acyl. Overall, the lipid can thus be classified as a structural subspecies.

```
      "fattyAcids": {
```

```
        "LCB": {
          "type": "STRUCTURAL",
          "name": "LCB",
          "position": 1,
          "lipidFaBondType": "ESTER",
          "lcb": true,
          "modifications": [],
          "doubleBondPositions": {},
          "ncarbon": 18,
          "nhydroxy": 2,
          "ndoubleBonds": 1
        },
        "FA1": {
          "type": "ISOMERIC",
          "name": "FA1",
          "position": 2,
          "lipidFaBondType": "ESTER",
          "lcb": false,
          "modifications": [],
          "doubleBondPositions": {
            "6": "Z"
          },
          "ncarbon": 16,
          "nhydroxy": 0,
          "ndoubleBonds": 1
        }
      }
```

Finally, the response reports the total number lipid names received, the total number parsed and the total number of parsing failures.

```
  ],
  "totalReceived": 1,
  "totalParsed": 1,
  "failedToParse": 0
}
```

### 2 C++ Implementation

This is the documentation for the Goslin reference implementation for C++. Please be aware, that the documentation is dedicated to developers of tools for computational lipidomics who want to use cppgoslin within their project. If you are interested to run Goslin as a user, please read Supplementary Section 1. The cppgoslin implementation has been developed with the following objectives:

1. To ease the handling with lipid names for developers working on mass spectrometry-based lipidomics tools.
2. To offer a tool that unifies all existing dialects of lipid names.

It is an open-source package under the MIT License available via github[1]. For a detailed structure of the implementation, read Supplementary Section 6.

#### Prerequisites

The cppgoslin library needs a GNU g++ compiler version with support for the C++ 11 standard. It comes with simple makefiles for easy compilation and installation. You need the following packages:

```
$ g++ (compiler)
$ make
```

To install the library globally on your system, simply type:

```
$ [sudo] make install
```

Be sure that you have root permissions. Here, the library and headers are installed into the /usr directory. If you want to change that location, you have to edit the first line within the *makefile*.

#### Testing cppgoslin

We set up more than 150 000 single unit and integration tests, to ensure that cppgoslin is parsing correctly. To run the tests, please type:

```
$ make test
$ make runtests
```

If a test should fail, please contact the developers[2].

[1] https://github.com/lifs-tools/cppgoslin
[2]

#### Using cppgoslin

The two major functions within cppgoslin are the parsing and printing of lipid names. A minimalistic example will demonstrate both functions the easiest way. In the examples folder, you will find the *lipid_name_parser.cpp* file. Compile it by typing:

```
$ cd examples
$ make
$ ./lipid_name_parser
```

Here is the minimalistic C++ code:

```
#include "cppgoslin/cppgoslin.h"
#include <iostream>
int main(){
    LipidParser parser;
    try {
        LipidAdduct* lipid = parser.parse("PA(12:0_14:0)");
        cout << lipid->get_lipid_string() << endl;
        delete lipid;
    }
    catch(LipidException& e){
        // handle the exception
        cout << e.what() << endl;
    }
    return 0;
}
```

To handle unexpected behavior, the parsing command should always be placed within a try/catch block and the `LipidAdduct` pointer should be deleted after usage to avoid memory leaks. Be aware when changing the installation directory, you also have to change the library directory within the examples *makefile*.

To retrieve a parsed lipid name on a higher hierarchy of lipid level, simply define the level when requesting the lipid name:

```
#include "cppgoslin/cppgoslin.h"
#include <iostream>
int main(){
    LipidParser parser;
    try {
        // providing a lipid name on isomeric subspecies level
        LipidAdduct* lipid = parser.parse("PA(12:1(5Z)/14:0)");
        cout << lipid->get_lipid_string(ISOMERIC_SUBSPECIES) << endl;
        cout << lipid->get_lipid_string(STRUCTURAL_SUBSPECIES) << endl;
        cout << lipid->get_lipid_string(MOLECULAR_SUBSPECIES) << endl;
        cout << lipid->get_lipid_string(SPECIES) << endl;
        cout << lipid->get_lipid_string(CLASS) << endl;
        cout << lipid->get_lipid_string(CATEGORY) << endl;
        delete lipid;
    }
    catch(LipidException& e){
        // handle the exception
        cout << e.what() << endl;
    }
    return 0;
}
```

Requesting a lipid name on a lower level than the provided will throw an exception. This functionality especially enables an easy way for computing data for histograms on lipid class or category level.

To increase the parsing performance, one can pick a parser for only one specific grammar:

```
GoslinParser goslin_parser;
GoslinFragmentParser goslin_fragment_parser;
LipidMapsParser lipid_maps_parser;
SwissLipidsParser swiss_lipids_parser;
HmdbParser hmdb_parser;
```

### 3 Python Implementation

This is the documentation for the Goslin reference implementation for Python 3. Please be aware, that the documentation is dedicated to developers of tools for computational lipidomics who want to insert pygoslin into their project. If you are interested to run Goslin as a user, please read Section 1. The pygoslin implementation has been developed with the following objectives:

1. To ease the handling with lipid names for developers working on mass spectrometrybased lipidomics tools.
2. To offer a tool that unifies all existing dialects of lipid names.

It is an open-source package under the MIT License available via github[3]. For a detailed structure of the implementation, read Supplementary Section 6.

#### Prerequisites

The pygoslin package uses Python's package management system *pip* to create an isolated and defined build environment. You need Python >=3.5 and the following packages to build the pygoslin package:

```
python3-pip
cython (module for Python 3)
make (optional)
```

To install the package globally in your Python distribution, simply type:

```
$ [sudo] make install
```

or

```
$ [sudo] python setup.py install
```

Be sure that you have root permissions.

#### Testing pygoslin

We set up more than 150 000 single unit and integration tests, to ensure that pygoslin is parsing correctly. To run the tests, please type:

```
$ make test
```

or

[3] https://github.com/lifs-tools/pygoslin

```
$ python3 -m unittest pygoslin.tests.FattyAcidTest
$ python3 -m unittest pygoslin.tests.ParserTest
$ python3 -m unittest pygoslin.tests.SwissLipidsTest
$ python3 -m unittest pygoslin.tests.GoslinTest
$ python3 -m unittest pygoslin.tests.LipidMapsTest
$ python3 -m unittest pygoslin.tests.HmdbTest
```

#### Using pygoslin

The two major functions within pygoslin are the parsing and printing of lipid names. You have several options, to access these functions. This example will demonstrate both functions the easiest way. Open a Python shell and type in:

```
from pygoslin.parser.Parser import LipidParser

lipid_parser = LipidParser() # setup the parser
lipid_name = "PE 16:1-12:0"

try:
    lipid = lipid_parser.parse(lipid_name) # start parsing
    print(lipid.get_lipid_string())
except Exception as e:
    print(e) # handle the exception
```

For all unexpected states, an exception is being raised. Be aware, that this method uses all available grammars in turn until a lipid name can be parsed successfully by a parser. Currently, five grammars are available, namely: Goslin, GoslinFragment, LipidMaps, SwissLipids, HMDB. To use a specific grammar / parser, you can use the following code:

```
# using solely the Goslin parser
from pygoslin.parser.Parser import GoslinParser
goslin_parser = GoslinParser()

lipid_name = "Cer 18:1;2/12:0"
try:
    lipid = goslin_parser.parse(lipid_name)
    print(lipid.get_lipid_string())
except Exception as e:
    print(e)
```

```
# using solely the Goslin Fragment parser
from pygoslin.parser.Parser import GoslinFragmentParser
goslin_fragment_parser = GoslinFragmentParser()

lipid_name = "Cer 18:1;2/12:0"
try:
    lipid = goslin_fragment_parser.parse(lipid_name)
    print(lipid.get_lipid_string())
except Exception as e:
    print(e)
```

```python
# using solely the LipidMaps parser
from pygoslin.parser.Parser import LipidMapsParser
lipid_maps_parser = LipidMapsParser()

lipid_name = "Cer(d18:1/12:0)"
try:
    lipid = lipid_maps_parser.parse(lipid_name)
    print(lipid.get_lipid_string())
except Exception as e:
    print(e)
```

```python
# using solely the SwissLipids parser
from pygoslin.parser.Parser import SwissLipidsParser
swiss_lipids_parser = SwissLipidsParser()

lipid_name = "Cer(d18:1/12:0)"
try:
    lipid = swiss_lipids_parser.parse(lipid_name)
    print(lipid.get_lipid_string())
except Exception as e:
    print(e)
```

```python
# using solely the HMDB parser
from pygoslin.parser.Parser import HmdbParser
hmdb_parser = HmdbParser()

lipid_name = "Cer(d18:1/12:0)"
try:
    lipid = hmdb_parser.parse(lipid_name)
    print(lipid.get_lipid_string())
except Exception as e:
    print(e)
```

To be as generic as possible, no treatment of validation of the fragment is conducted within the GoslinFragmentParser.

To retrieve a parsed lipid name on a higher hierarchy of lipid level, simply define the level when requesting the lipid name:

```python
# report on different lipid hierarchies
from pygoslin.parser.Parser import *
from pygoslin.domain.LipidLevel import LipidLevel

parser = LipidParser()
# providing a lipid name on isomeric subspecies level
lipid_name = "PA 18:1(5Z)/12:0"
```

```python
try:
    lipid = parser.parse(lipid_name)
    print(lipid.get_lipid_string(LipidLevel.ISOMERIC_SUBSPECIES))
    print(lipid.get_lipid_string(LipidLevel.STRUCTURAL_SUBSPECIES))
    print(lipid.get_lipid_string(LipidLevel.MOLECULAR_SUBSPECIES))
    print(lipid.get_lipid_string(LipidLevel.SPECIES))
    print(lipid.get_lipid_string(LipidLevel.CLASS))
    print(lipid.get_lipid_string(LipidLevel.CATEGORY))
except Exception as e:
    print(e)
```

This functionality especially enables an easy way for computing data for histograms on lipid class or category level. Requesting a lipid name on a lower level than the provided will raise an exception.

### 4 R Implementation

This project is a parser, validator and normalizer implementation for shorthand lipid nomenclatures, using the Grammar of Succinct Lipid Nomenclatures project for the R language [4].

Goslin defines multiple grammars compatible with ANTLRv4 for different sources of shorthand lipid nomenclature. This allows to generate parsers based on the defined grammars, which provide immediate feedback whether a processed lipid shorthand notation string is compliant with a particular grammar, or not.

rgoslin uses the Goslin grammars and the cppgoslin parser to support the following general tasks:

1. Facilitate the parsing of shorthand lipid names dialects.
2. Provide a structural representation of the shorthand lipid after parsing.
3. Use the structural representation to generate normalized names.

rgoslin is an open-source package available via github[5].

#### Prerequisites

This project uses the R programming language. To be able to use it, please install R[6] following the instructions for your particular operating system. rgoslin is based on native C++ code (via cppgoslin). It therefore requires additional tools on your system to compile and install it. Please see the Rcpp FAQ[7], question 1.3 for installation details for your specific operating system.

Install the ‘devtools‘ package with the following command.

```
if(!require(devtools)) { install.packages("devtools") }
```

Run

```
install_github("lifs-tools/rgoslin")
```

to install from the github repository.

This will install the latest, potentially unstable development version of the package with all required dependencies into your local R installation.

If you want to use a proper release version, referenced by a Git tag (here: v1.0.0) install the package as follows:

[4] https://www.r-project.org/
[5] https://github.com/lifs-tools/rgoslin
[6] https://cloud.r-project.org/
[7] https://cran.r-project.org/web/packages/Rcpp/vignettes/Rcpp-FAQ.pdf

```
install_github("lifs-tools/rgoslin", ref="v1.0.0")
```

If you have cloned the code locally, use devtools as follows. Make sure you set the working directory to where the API code is located. Then execute

```
library(devtools)
install(".")
```

#### Testing rgoslin

rgoslin uses the testthat R package to provide unit tests for the lipid name parsing methods. The tests are located in the `tests` folder. To run the tests, execute

```
library(devtools)
test()
```

#### Using rgoslin

To load the package, start an R session and type

```
library(rgoslin)
```

Type the following to see the package vignette / tutorial:

```
vignette('introduction', package = 'rgoslin')
```

In order to use the provided translation functions of rgoslin, you first need to load the library.

```
library(rgoslin)
```

To check, whether a given lipid name can be parsed by any of the parsers supplied by cppgoslin, you can use the `isValidLipidName` method. It will return `TRUE` if the given name can be parsed by any of the available parsers and `FALSE` if the name was not parseable.

```
isValidLipidName("PC 32:1")
```

Using `parseLipidName` with a lipid name returns a named vector of properties of the parsed lipid name.

```
pc32vector <- parseLipidName("PC 32:1")
pc32df <- as.data.frame(t(pc32vector))
```

If you want to set the grammar to parse against manually, this is also possible:

```
originalName <- "TG(16:1(5E)/18:0/20:2(3Z,6Z))"
tagVec <- rgoslin::parseLipidNameWithGrammar(originalName, "LipidMaps")
tagDf <- as.data.frame(t(tagVec))
```

Currently, the following grammars are available: LipidMaps, SwissLipids, Goslin, Goslin-Fragments, HMDB.

If you want to parse multiple lipid names, use the `parseLipidNames` method with a vector of lipid names. This returns a data frame of properties of the parsed lipid names with one row per lipid.

```
multipleLipidNames <- parseLipidNames(c("PC 32:1","LPC 34:1","TG(18:1_18:0_16:1)"))
```

Finally, if you want to parse multiple lipid names and want to use one particular grammar:

```
originalNames <- c("PC 32:1","LPC 34:1","TAG 18:1_18:0_16:1")
multipleLipidNamesWithGrammar <- parseLipidNamesWithGrammar(originalNames,
    "Goslin")
```

### 5 Java Implementation

This project is a parser, validator and normalizer implementation for shorthand lipid nomenclatures, based on Goslin for the Java programming language[8].

Goslin defines multiple grammars compatible with ANTLRv4 for different sources of shorthand lipid nomenclature. This allows to generate parsers based on the defined grammars, which provide immediate feedback whether a processed lipid shorthand notation string is compliant with a particular grammar, or not.

Here, jgoslin uses the Goslin grammars and the generated parsers to support the following general tasks:

1. Facilitate the parsing of shorthand lipid names dialects.
2. Provide a structural representation of the shorthand lipid after parsing.
3. Use the structural representation to generate normalized names.

Furthermore, jgoslin is an open-source package available via github[9].

##### Prerequisites

This project is based on Java 11. To use it, you need a Java Runtime Environment (JRE) installed on your system. If you want to use the library in your own Java projects, you need a Java Development Kit (JDK) installed on your system. Please consult `https://adoptopenjdk.net/installation.html` for installation options and instructions for your operating system.

Installation instructions

Building the project and generating client code from the command-line

In order to build the client code and run the unit tests, execute the following command from a terminal:

```
./mvnw install
```

or on Windows:

```
mvnw.bat install
```

This compiles and tests the Java library.

[8] https://go.java/

[9] https://github.com/lifs-tools/jgoslin

#### Testing jgoslin

Here, jgoslin comes with a comprehensive collection of unit (JUnit 5), integration (JUnit 5) and acceptance (Cucumber) tests. You can run all of them as follows:

```
./mvnw verify
```

#### Using the command-line interface

The `cli` sub-project provides a command line interface (CLI) for parsing of lipid names either from the command line or from a file with one lipid name per line.

After building the project as mentioned above with `./mvnw install`, the `cli/target` folder will contain the `jgoslin-cli-<VERSION>-bin.zip` file. Alternatively, you can download the latest cli zip file from Bintray: https://bintray.com/lifs/maven/jgoslin-cli[Search for latest jgoslin-cli-<VERSION>-bin.zip artefact] and click to download.

In order to run the validator, unzip that file, change into the unzipped folder and run

```
java -jar jgoslin-cli-<VERSION>.jar
```

to see the available options.

To parse a single lipid name from the command line using all available parsers, run

```
java -jar jgoslin-cli-<VERSION>.jar -n "Cer(d18:1/20:2)"
```

The output will tell you what is done and will echo a table of the results to the terminal:

```
Parsing lipid identifier: Cer(d18:1/20:2)
Parsing lipid identifier: Cer(d18:1/20:2)
Parsing lipid maps identifier: Cer(d18:1/20:2)
Parsing swiss lipids identifier: Cer(d18:1/20:2)
Parsing HMDB lipids identifier: Cer(d18:1/20:2)
Echoing output to stdout.
Normalized Name Original Name Grammar Message Lipid Maps Category Lipid Maps Main
    Class Functional Class Abbr Functional Class Synonyms Level Total #C Total #OH
    Total #DB LCB SN Position LCB #C LCB #OH LCB #DB LCB Bond Type FA1 SN Position
    FA1 #C FA1 #OH FA1 #DB FA1 Bond Type
Cer(d18:1/20:2) GOSLIN no viable alternative at input 'Cer('
Cer(d18:1/20:2) GOSLIN_FRAGMENTS no viable alternative at input 'Cer('

Cer 18:1;2/20:2 Cer(d18:1/20:2) LIPIDMAPS  Sphingolipid [SP]
    N-acyl-4-hydroxysphinganines (phytoceramides) [SP0203] [Cer] [Cer, Ceramide]
    STRUCTURAL_SUBSPECIES 38 2 3 1 18 2 1 ESTER 2 20 0 2 ESTER
Cer 18:1;2/20:2 Cer(d18:1/20:2) SWISSLIPIDS  Sphingolipid [SP]
    N-acyl-4-hydroxysphinganines (phytoceramides) [SP0203] [Cer] [Cer, Ceramide]
    STRUCTURAL_SUBSPECIES 38 2 3 1 18 2 1 ESTER 2 20 0 2 ESTER
Cer 18:1;2/20:2 Cer(d18:1/20:2) HMDB  Sphingolipid [SP]
    N-acyl-4-hydroxysphinganines (phytoceramides) [SP0203] [Cer] [Cer, Ceramide]
    STRUCTURAL_SUBSPECIES 38 2 3 1 1821 ESTER 2 20 0 2 ESTER
```

To parse multiple lipid names from a file via the commmand line, run

```
java -jar jgoslin-cli-<VERSION>.jar -f lipidNames.txt
```

To use a specific grammar, instead of trying all, run

```
java -jar jgoslin-cli-<VERSION>.jar -f lipidNames.txt -g GOSLIN
```

To write output to the tab-separated output file 'goslin-out.tsv' instead of to the terminal, run

```
java -jar jgoslin-cli-<VERSION>.jar -f lipidNames.txt -g GOSLIN -o
```

If you want to use all available grammars, simply omit the `-g GOSLIN` argument. Please note that you will then receive N times M lines in the output file, where N is the number of lipid names and M the number of grammars.

#### Using jgoslin

To integrate jgoslin in your own projects as a library, please see the README file at `https://github.com/lifs-tools/jgoslin` for more details.

The following snippet shows how to parse a shorthand lipid name with the different parsers:

```
import de.isas.lipidomics.domain.*; // contains Domain objects like LipidAdduct,
    LipidSpecies ...
import de.isas.lipidomics.palinom.*; // contains the parser implementations
...

String ref = "Cer(d18:1/20:2)";
try {
   // use the SwissLipids parser
   SwissLipidsVisitorParser slParser = new SwissLipidsVisitorParser();
   LipidAdduct sllipid = slParser.parse(ref);
   System.out.println(sllipid.getLipidString()); // to print the lipid name to
       the console
} catch (ParsingException pe) {
// catch this for any syntactical issues with the name during parsing with a
    particular parser
   pe.printStackTrace();
} catch (ParseTreeVisitorException ptve) {
// catch this for any structural issues with the name during parsing with a
    particular parser
   ptve.printStackTrace();
}

//alternatively, use the other parsers. Don't forget to place try catch blocks
    around the following lines, as for the SwissLipids parser example
   // use the LipidMAPS parser
   LipidMapsVisitorParser lmParser = new LipidMapsVisitorParser();
   LipidAdduct lmlipid = lmParser.parse(ref);
   // use the shorthand notation parser GOSLIN
   GoslinVisitorParser goslinParser = new GoslinVisitorParser();
   LipidAdduct golipid = goslinParser.parse(ref);
   // use the shorthand notation parser with support for fragments
       GOSLIN_FRAGMENTS
```

```java
GoslinFragmentsVisitorParser goslinFragmentsParser = new
    GoslinFragmentsVisitorParser();
LipidAdduct gflipid = goslinFragmentsParser.parse(ref);
```

To retrieve a parsed lipid name on a higher hierarchy of lipid level, simply define the level when requesting the lipid name:

```java
System.out.println(sllipid.getLipidString(LipidLevel.CATEGORY));
System.out.println(sllipid.getLipidString(LipidLevel.CLASS));
System.out.println(sllipid.getLipidString(LipidLevel.SPECIES));
System.out.println(sllipid.getLipidString(LipidLevel.MOLECULAR_SUBSPECIES));
System.out.println(sllipid.getLipidString(LipidLevel.STRUCTURAL_SUBSPECIES));
System.out.println(sllipid.getLipidString(LipidLevel.ISOMERIC_SUBSPECIES)); //
    will throw a ConstraintViolationException since this lipid is only on
    structural subspecies level
```

This functionality allows easy computation of aggregate statistics of lipids on lipid class, category or arbitrary levels. Requesting a lipid name on a lower level than the provided will raise an exception.

For an overview of the domain model used by jgoslin, please see Supplementary Section 6.

### 6 Goslin Object Model

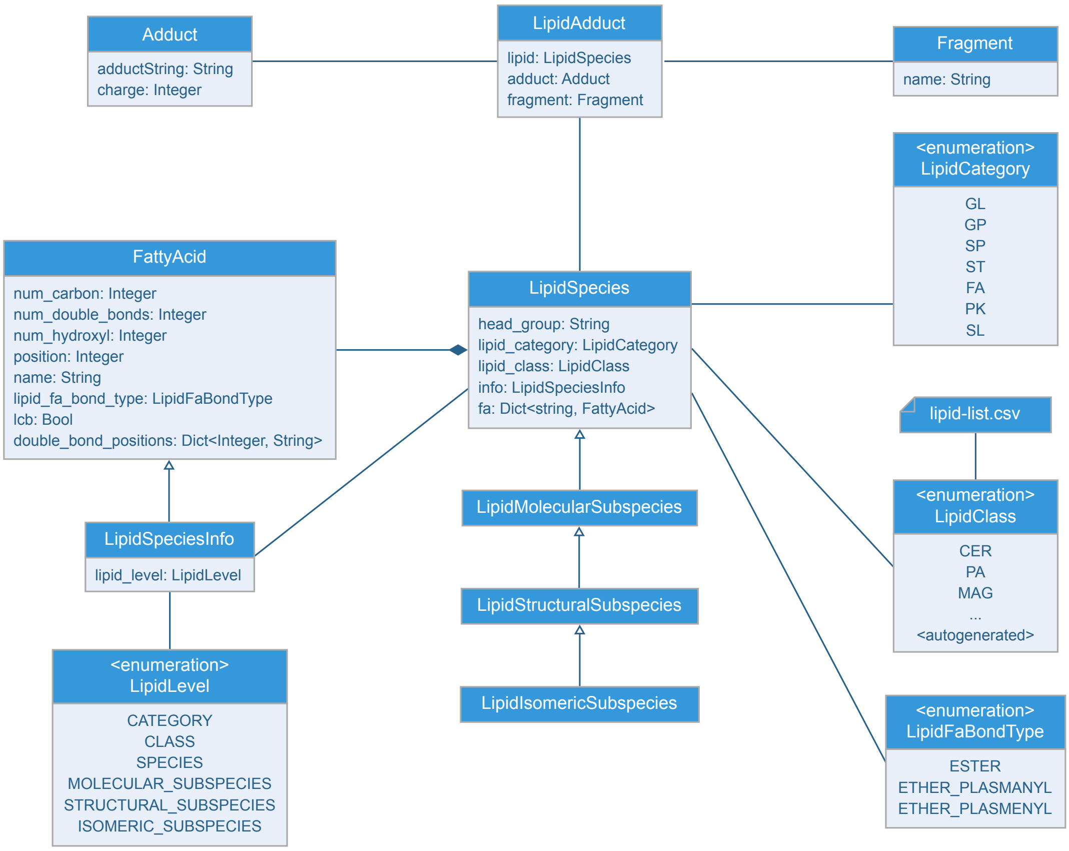


Supplementary Figure 7: Goslin object model.

All goslin implementations are implementing the goslin object model as illustrated in Supplementary Figure 7. The classes `LipidCategory`, `LipidLevel`, `LipidClass`, and `LipidFaBondType` are predefined enumerations. Here, `LipidClass` is being generated automatically from a list containing lipid information (name, description, category, abbreviation, synonyms) for all implementations, see Supplementary Table 1 for details. This especially eases the maintenance and ensures that the goslin implementations have the same data base. The main class unifying all classes and being provided by the parsers is `LipidAdduct`. It contains information about the pure lipid, the adduct as well as the fragment (if defined). The different lipid classes inherit from each other in a hierarchical fashion as defined by Liebisch et al.[4]. A dictionary with the class `LipidSpecies` is storing all its associated fatty acyl chains which are defined within the class `FattyAcid`. For storing the cummulated information on species level for the carbon length, double bonds, etc, the class `LipidSpeciesInfo` is utilized.

### 7 List of Supported Lipids

Supplementary Table 1: List of supported lipids, lipid classes and their normalized abbreviations

| Category | Description | Abbreviation |
|---|---|---|
| Fatty acyls | Other Docosanoids | 10-HDoHE |
| | Epoxyeicosatrienoic acids | 11(12)-EET |
| | Hydroxy/hydroperoxyeicosatetraenoic acids | 11,12-DHET |
| | Other Docosanoids | 11-HDoHE |
| | Hydroxy/hydroperoxyeicosatetraenoic acids | 11-HETE |
| | Other Octadecanoids | 12(13)-EpOME |
| | Hydroxy/hydroperoxyeicosapentaenoic acids | 12-HEPE |
| | Hydroxy/hydroperoxyeicosatetraenoic acids | 12-HETE |
| | Hydroxy/hydroperoxyeicosatrienoic acids | 12-HHTrE |
| | Fatty acids and conjugates | 12-OxoETE |
| | Other Octadecanoids | 13-HODE |
| | Other Octadecanoids | 13-HOTrE |
| | Epoxyeicosatrienoic acids | 14(15)-EET |
| | Other Eicosanoids | 14(15)-EpETE |
| | Hydroxy/hydroperoxyeicosatetraenoic acids | 14,15-DHET |
| | Hydroxy/hydroperoxyeicosapentaenoic acids | 15-HEPE |
| | Hydroxy/hydroperoxyeicosatetraenoic acids | 15-HETE |
| | Prostaglandins | 15d-PGJ2 |
| | Other Docosanoids | 16-HDoHE |
| | Hydroxy/hydroperoxyeicosatetraenoic acids | 16-HETE |
| | Hydroxy/hydroperoxyeicosapentaenoic acids | 18-HEPE |
| | Epoxyeicosatrienoic acids | 5(6)-EET |
| | Hydroxy/hydroperoxyeicosatetraenoic acids | 5,12-DiHETE |
| | Lipoxins | 5,6,15-LXA4 |
| | Hydroxy/hydroperoxyeicosatetraenoic acids | 5,6-DiHETE |
| | Hydroxy/hydroperoxyeicosapentaenoic acids | 5-HEPE |
| | Hydroxy/hydroperoxyeicosatetraenoic acids | 5-HETE |
| | Hydroxy/hydroperoxyeicosatetraenoic acids | 5-HpETE |
| | Fatty acids and conjugates | 5-OxoETE |
| | Epoxyeicosatrienoic acids | 8(9)-EET |
| | Hydroxy/hydroperoxyeicosatetraenoic acids | 8,9-DHET |
| | Other Docosanoids | 8-HDoHE |
| | Hydroxy/hydroperoxyeicosatetraenoic acids | 8-HETE |
| | Other Octadecanoids | 9(10)-EpOME |
| | Hydroxy/hydroperoxyeicosapentaenoic acids | 9-HEPE |
| | Hydroxy/hydroperoxyeicosatetraenoic acids | 9-HETE |
| | Other Octadecanoids | 9-HODE |
| | Other Octadecanoids | 9-HOTrE |
| | Unsaturated fatty acids | AA |
| | Fatty acyl carnitines | CAR |
| | Fatty acyl CoAs | CoA |

| Category | Description | Abbreviation |
|---|---|---|
| Fatty acyls | Unsaturated fatty acids | DHA |
| | Unsaturated fatty acids | EPA |
| | Fatty acids and conjugates | FA |
| | Fatty acyl | FA |
| | Wax monoesters | FAHFA |
| | Glycerophosphoethanolamine | GP-NAE |
| | Leukotrienes | LTB4 |
| | Eicosanoid derivatives | LTC4 |
| | Leukotrienes | LTD4 |
| | Unsaturated fatty acids | Linoleic acid |
| | Maresins | Maresin 1 |
| | Fatty amides | NAE |
| | Prostaglandins | PGB2 |
| | Prostaglandins | PGD2 |
| | Prostaglandins | PGE2 |
| | Prostaglandins | PGF2alpha |
| | Prostaglandins | PGI2 |
| | Straight chain fatty acids | Palmitic acid |
| | Resolvin Ds | Resolvin D1 |
| | Resolvin Ds | Resolvin D2 |
| | Resolvin Ds | Resolvin D3 |
| | Resolvin Ds | Resolvin D5 |
| | Thromboxanes | TXB1 |
| | Thromboxanes | TXB2 |
| | Thromboxanes | TXB3 |
| | Fatty esters | WE |
| | Fatty acids and conjugates | alpha-LA |
| | Hydroxy/hydroperoxyeicosatetraenoic acids | tetranor-12-HETE |
| Glycero-lipids | Diacylglycerols | DAG |
| | Other Glycerolipids | DGCC |
| | Glycosyldiradylglycerols | DGDG |
| | Dihexosyldiacylglycerol | DHDG |
| | Monoacylglycerols | MAG |
| | Glycosyldiacylglycerols | MGDG |
| | Monohexosyldiacylglycerol | MHDG |
| | Glycosyldiradylglycerols | SQDG |
| | Glycosylmonoacylglycerols | SQMG |
| | Triacylglycerols | TAG |
| Glycero-phospho-lipids | Glycosylglycerophospholipids | 6-Ac-Glc-GP |
| | Monoacylglycerophosphomonoradylglycerols | BMP |
| | CDP-diacylglycerols | CDPDAG |
| | Cardiolipins | CL |
| | Glycerophosphoinositolglycans | CPA |
| | Glycerophosphoglycerophosphoglycerols | DLCL |
| | Dimethylphosphatidylethanolamine | DMPE |
| | Glycosyldiradylglycerols | Glc-DG |

| Category | Description | Abbreviation |
|---|---|---|
| | Diacylglycosylglycerophospholipids | Glc-GP |
| | Lyso-CDP-diacylglycerol | LCDPDAG |
| | Lysodimethylphosphatidylethanolamine | LDMPE |
| | Lysomonomethylphosphatidylethanolamine | LMMPE |
| | Monoacylglycerophosphates | LPA |
| | Monoacylglycerophosphocholines | LPC |
| | Monoacylglycerophosphoethanolamines | LPE |
| | 1Z-alkenylglycerophosphoglycerols | LPG |
| | Monoacylglycerophosphoinositols | LPI |
| | Monoacylglycerophosphoinositolglycans | LPIM1 |
| | Glycerophosphoinositolglycans | LPIM2 |
| | Glycerophosphoinositolglycans | LPIM3 |
| | Glycerophosphoinositolglycans | LPIM4 |
| | Glycerophosphoinositolglycans | LPIM5 |
| | Glycerophosphoinositolglycans | LPIM6 |
| | Lysophosphatidylinositol- mannosideinositolphosphate | LPIMIP |
| | Lysophosphatidylinositol-glucosamine | LPIN |
| | Monoacylglycerophosphoserines | LPS |
| | Glycerophosphoglycerophosphoglycerols | MLCL |
| | Monomethylphosphatidylethanolamine | MMPE |
| Glycero-phospho-lipids | Glycerophosphoethanolamine | NAPE |
| | Diacylglycerophosphates | PA |
| | Oxidized glycerophosphocholines | PC |
| | Oxidized glycerophosphoethanolamines | PE |
| | Glycerophosphoethanolamines | PE-NMe |
| | Glycerophosphoethanolamines | PE-NMe2 |
| | Glycerophosphoethanolamines | PEt |
| | Diacylglycerophosphoglycerols | PG |
| | Diacylglycerophosphoglycerophosphates | PGP |
| | Diacylglycerophosphoinositols | PI |
| | Diacylglycerophosphoinositolglycans | PIM1 |
| | Glycerophosphoinositolglycans | PIM2 |
| | Glycerophosphoinositolglycans | PIM3 |
| | Glycerophosphoinositolglycans | PIM4 |
| | Glycerophosphoinositolglycans | PIM5 |
| | Glycerophosphoinositolglycans | PIM6 |
| | Phosphatidylinositol mannoside inositol phosphate | PIMIP |
| | Diacylglycerophosphoinositol monophosphates | PIP |
| | Diacylglycerophosphoinositol bisphosphates | PIP2 |
| | Glycerophosphoinositolbisphosphates | PIP2[3',4'] |
| | Glycerophosphoinositolbisphosphates | PIP2[3',5'] |
| | Glycerophosphoinositolbisphosphates | PIP2[4',5'] |
| | Diacylglycerophosphoinositol trisphosphates | PIP3 |
| | Glycerophosphoinositoltrisphosphates | PIP3[3',4',5'] |
| | Glycerophosphoinositolmonophosphates | PIP[3'] |

| Category | Description | Abbreviation |
| --- | --- | --- |
| | Glycerophosphoinositolmonophosphates | PIP[4'] |
| | Glycerophosphoinositolmonophosphates | PIP[5'] |
| | Diacylglyceropyrophosphates | PPA |
| | Diacylglycerophosphoserines | PS |
| | Diacylglycerophosphoserines | PS-NAc |
| | Other Glycerophospholipids | PT |
| | Glycerophosphonocholines | PnC |
| | Glycerophosphoinositolglycans | PnE |
| | Diacylglycerophosphomonoradylglycerols | SLBPA |
| Saccharo-lipids | Acyltrehaloses | AC2SGL |
| | Acyltrehaloses | DAT |
| | Acyltrehaloses | PAT16 |
| | Acyltrehaloses | PAT18 |
| Sphingo-lipids | Glycosphingolipids | (3'-sulfo)LacCer |
| | Glycosphingolipids | (Fuc)iGb3Cer |
| | Acylceramides | 1-O-behenoyl-Cer |
| | Acylceramides | 1-O-carboceroyl-Cer |
| | Acylceramides | 1-O-cerotoyl-Cer |
| | Acylceramides | 1-O-eicosanoyl-Cer |
| | Acylceramides | 1-O-lignoceroyl-Cer |
| | Acylceramides | 1-O-myristoyl-Cer |
| | Acylceramides | 1-O-palmitoyl-Cer |
| | Acylceramides | 1-O-stearoyl-Cer |
| | Acylceramides | 1-O-tricosanoyl-Cer |
| | Globoside | Ac-O-9-GD1a |
| | Globoside | Ac-O-9-GT1b |
| | Globoside | Ac-O-9-GT3 |
| | Glycosphingolipids | Branched-Forssman |
| | Ceramide-1-phosphates | C1P |
| | N-acylsphingosines (ceramides) | Cer |
| | Ceramide 1-phosphates | CerP |
| | Glycosphingolipids | DSGG |
| | Ceramide phosphoethanolamines | EPC |
| | Simple Glc series | FMC-5 |
| | Neutral glycosphingolipids | FMC-6 |
| | Glycosphingolipids | Forssman |
| | Acidic glycosphingolipids | Fuc(Gal)-GM1 |
| | Glycosphingolipids | Fuc(Gal)Gal-iGb4Cer |
| | Glycosphingolipids | Fuc-Branched-Forssman |
| | Globoside | Fuc-GA1 |
| | Globoside | Fuc-GD1b |
| | Globoside | Fuc-GM1 |
| | Globoside | Fuc-GM1(NeuGc) |
| | Glycosphingolipids | Fuc-iGb3Cer |

| Category | Description | Abbreviation |
|---|---|---|
| Sphingo-lipids | Glycosphingolipids | FucGalGb3Cer |
| | Glycosphingolipids | GA1 |
| | Glycosphingolipids | GA2 |
| | Neutral glycosphingolipids | GB4 |
| | Glycosphingolipids | GD1 |
| | Ganglioside GD1a(d18:1(4E)) | GD1a |
| | Ganglioside GD1a alpha(d18:1(4E)) | GD1a alpha |
| | Globoside | GD1a(NeuAc/NeuGc) |
| | Globoside | GD1a(NeuGc/NeuAc) |
| | Globoside | GD1a(NeuGc/NeuGc) |
| | Ganglioside GD1b(d18:1(4E)) | GD1b |
| | Ganglioside GD1c(d18:1(4E)) | GD1c |
| | Globoside | GD1c(NeuGc/NeuGc) |
| | Glycosphingolipids | GD2 |
| | Glycosphingolipids | GD3 |
| | Glycosphingolipids | GM1 |
| | Globoside | GM1 alpha |
| | Globoside | GM1(NeuGc) |
| | Ganglioside GM1b(d18:1(4E)) | GM1b |
| | Globoside | GM1b(NeuGc) |
| | Glycosphingolipids | GM2 |
| | Globoside | GM2(NeuGc) |
| | Glycosphingolipids | GM3 |
| | Gangliosides | GM4 |
| | Glycosphingolipids | GP1 |
| | Ganglioside GP1c(d18:1(4E)) | GP1c |
| | Ganglioside GP1c alpha(d18:1(4E)) | GP1c alpha |
| | Glycosphingolipids | GQ1 |
| | Ganglioside GQ1b(d18:1(4E)) | GQ1b |
| | Ganglioside GQ1b alpha(d18:1(4E)) | GQ1b alpha |
| | Ganglioside GQ1c(d18:1(4E)) | GQ1c |
| | Glycosphingolipids | GT1 |
| | Ganglioside GT1a(d18:1(4E)) | GT1a |
| | Ganglioside GT1a alpha(d18:1(4E)) | GT1a alpha |
| | Ganglioside GT1b(d18:1(4E)) | GT1b |
| | Globoside | GT1b alpha |
| | Globoside | GT1b alpha(NeuGc) |
| | Ganglioside GT1c(d18:1(4E)) | GT1c |
| | Glycosphingolipids | GT2 |
| | Glycosphingolipids | GT3 |
| | Globoside | Gal(Fuc)-GA1 |
| | Globoside | Gal(Fuc)-GD1b |
| | Globoside | Gal-GD1b |
| | Glycosphingolipids | Gal-iGb4Cer |
| | Globoside | GalGal-GD1b |
| | Glycosphingolipids | GalGalGalGb3Cer |

| Category | Description | Abbreviation |
|---|---|---|
| Sphingo-lipids | Glycosphingolipids | GalGalGb3Cer |
| | Globoside | GalGalNAc-GM1b(NeuGc) |
| | Glycosphingolipids | GalGb3Cer |
| | Glycosphingolipids | GalGb4Cer |
| | Glycosphingolipids | GalGlcNAc-GalGb4Cer |
| | Globoside | GalNAc-GD1a |
| | Globoside | GalNAc-GD1a(NeuAc/NeuGc) |
| | Globoside | GalNAc-GD1a(NeuGc/NeuAc) |
| | Globoside | GalNAc-GM1 |
| | Globoside | GalNAc-GM1b |
| | Globoside | GalNAc-GM1b(NeuGc) |
| | Globoside | GalNAcGal(Fuc)-GA1 |
| | Glycosphingolipids | GalNAcGalGb3Cer |
| | Glycosphingolipids | Gb3 |
| | Glycosphingolipids | Gb3Cer |
| | Glycosphingolipids | Gb4Cer |
| | Glycosphingolipids | GlcNAc-GalGb4Cer |
| | Glycosphingolipids | GlcNAcGb3Cer |
| | Glycosphingolipids | Globo-A |
| | Glycosphingolipids | Globo-B |
| | Glycosphingolipids | Globo-H |
| | Glycosphingolipids | Globo-Lex-9 |
| | Glycosphingolipids | Hex2Cer |
| | Neutral glycosphingolipids | Hex3Cer |
| | Glycosphingolipids | HexCer |
| | Ceramide phosphoinositols | IPC |
| | Sphinganines | LCB |
| | Sphingoid base 1-phosphates | LCBP |
| | Hexosylsphingosine | LHexCer |
| | Ceramides | LSM |
| | Globoside | Lex-GM1 |
| | Phosphosphingolipids | M(IP)2C |
| | Phosphosphingolipids | MIPC |
| | Glycosphingolipids | MSGG |
| | Glycosphingolipids | NOR1 |
| | Glycosphingolipids | NOR2 |
| | Glycosphingolipids | NORint |
| | Glycosphingolipids | NeuAc(alpha2-6)-MSGG |
| | Glycosphingolipids | NeuAc(alpha2-8)-MSGG |

| Category | Description | Abbreviation |
|---|---|---|
| Sphingo-lipids | Glycosphingolipids | NeuAcGal-iGb4Cer |
| | Glycosphingolipids | NeuGc-GalGb4Cer |
| | Globoside | NeuGc-LacNAc-GM1(NeuGc) |
| | Glycosphingolipids | NeuGcNeuGc-GalGb4Cer |
| | Glycosphingolipids | Para-Forssman |
| | Globoside | SB1a |
| | Glycosphingolipids | SHex2Cer |
| | Sulfoglycosphingolipids (sulfatides) | SHexCer |
| | Ceramide phosphocholines (sphingomyelins) | SM |
| | Globoside | SM1a |
| | Globoside | SM1b |
| | Globoside | SO3-GM1(NeuGc) |
| | Glycosphingolipids | SO3-Gal-iGb4Cer |
| | Glycosphingolipids | SO3-GalGb4Cer |
| | Glycosphingolipids | SO3-Gb4Cer |
| | Glycosphingolipids | SO3-iGb4Cer |
| | Glycosphingolipids | SulfoGalCer |
| | Glycosphingolipids | i-Forssman |
| | Glycosphingolipids | iGb3Cer |
| | Glycosphingolipids | iGb4Cer |
| Sterols | Sterol esters | SE |
| | Steryl esters | SE 27:1 |
| | Desmosterol Ester | SE 27:2 |
| | Ergostadienol Ester | SE 28:2 |
| | Ergosterol Ester | SE 28:3 |
| | Stigmasterol Ester | SE 29:2 |
| | Lanosterol Ester | SE 30:2 |
| | Sterols | ST |
| | Cholesterol and derivatives | ST 27:1;1 |
| | Desmosterol | ST 27:2;1 |
| | Ergostadienol | ST 28:2;1 |
| | Ergosterol | ST 28:3;1 |
| | Stigmasterol | ST 29:2;1 |
| | Lanosterol | ST 30:2;1 |
| Polyketides | Anacardic acids and derivatives | ANACARD |
| | Alkyl catechols and derivatives | CATECHOL |
| | Alkyl phenols and derivatives | PHENOL |
| | Alkyl resorcinols and derivatives | RESORCINOL |

#### Acronyms

**API** application programming interface. 6, 7

**CLI** command line interface. 20

**Goslin** grammar of succinct lipid nomenclatures. 2, 3, 6, 9, 12, 16, 19, 23

**HTTP** hypertext transfer protocol. 6, 7

**JDK** Java Development Kit. 19

**JRE** Java Runtime Environment. 19

**JSON** JavaScript object notation. 6

**LCB** long chain base. 7

**REST** representational state transfer. 6, 7

#### References


[1] Eoin Fahy, Shankar Subramaniam, Robert C. Murphy, Masahiro Nishijima, Christian R. H. Raetz, Takao Shimizu, Friedrich Spener, Gerrit van Meer, Michael J. O. Wakelam, and Edward A. Dennis. Update of the LIPID MAPS comprehensive classification system for lipids. *Journal of Lipid Research*, 50(Supplement):S9–S14, 2009.

[2] Lucila Aimo, Robin Liechti, Nevila Hyka-Nouspikel, Anne Niknejad, Anne Gleizes, Lou Götz, Dmitry Kuznetsov, Fabrice P.A. David, F. Gisou van der Goot, Howard Riezman, Lydie Bougueleret, Ioannis Xenarios, and Alan Bridge. The SwissLipids knowledgebase for lipid biology. *Bioinformatics*, 31(17):2860–2866, 2015.

[3] David S. Wishart, Dan Tzur, Craig Knox, Roman Eisner, An Chi Guo, Nelson Young, Dean Cheng, Kevin Jewell, David Arndt, Summit Sawhney, Chris Fung, Lisa Nikolai, Mike Lewis, Marie-Aude Coutouly, Ian Forsythe, Peter Tang, Savita Shrivastava, Kevin Jeroncic, Paul Stothard, Godwin Amegbey, David Block, David. D. Hau, James Wagner, Jessica Miniaci, Melisa Clements, Mulu Gebremedhin, Natalie Guo, Ying Zhang, Gavin E. Duggan, Glen D. MacInnis, Alim M. Weljie, Reza Dowlatabadi, Fiona Bamforth, Derrick Clive, Russ Greiner, Liang Li, Tom Marrie, Brian D. Sykes, Hans J. Vogel, and Lori Querengesser. HMDB: the Human Metabolome Database. *Nucleic Acids Research*, 35(suppl_1):D521–D526, 2007.

[4] Gerhard Liebisch, Juan Antonio Vizcaíno, Harald Köfeler, Martin Trötzmüller, William J. Griffiths, Gerd Schmitz, Friedrich Spener, and Michael J. O. Wakelam. Shorthand notation for lipid structures derived from mass spectrometry. *Journal of Lipid Research*, 54(6):1523–1530, 2013.
